# Multifrequency control of Faraday wave bioassembly for constructing multiscale hPSC-derived neuronal networks

**DOI:** 10.1101/2023.12.01.569533

**Authors:** Longjun Gu, Wen Zhao, Yuhang Fan, Jia Shang, Yang Zhao, Jibo Wang, Tao Chen, Peidi Liu, Pu Chen

## Abstract

Bioassembly is recently regarded as a critical alternative biofabrication technical route to bioprinting since it can directly manipulate millions of live cells to form multicellular structures with close intercellular proximity, improving contact-dependent cell communication and promoting the emergence of tissue-specific functions. However, acoustic bioassembly techniques are currently limited to generating cytoarchitecture with a single characteristic length which cannot faithfully mimic the multiscale cellular structures in native tissues. To overcome this challenge, herein we report a novel acoustic bioassembly technique that employs multifrequency control of Faraday waves to form multiscale cellular structures. By superimposing multiple sine wave signals with proper amplitude ratios, Faraday waves containing multiple wavelengths can be induced and enabled to generate multiscale structures in few seconds. Using this technique, we construct functional neuronal networks with multiscale connectivity that display spontaneous neuroelectrical activities. We anticipate this technique will find wide applications in tissue engineering and regenerative medicine.

## 1. Introduction

In the human body, the cytoarchitectures in tissues are usually highly complex and contain multiscale structures. The structural organization plays an important role in mimicking native cytoarchitectures and reconstituting tissue-specific physiological functions^[1–3]^. For example, the hepatic lobule is the liver’s basic functional unit and performs the most critical functions such as plasma protein synthesis, carbohydrate metabolism, and bile production. The hepatic lobule has a multiscale cytoarchitecture^[4]^. Specifically, the cross-sectional diameter of an independent hepatic lobule is almost 1 mm, while a radially arranged hepatic sinusoid is approximately 10 μm in diameter. As another example, the cerebral cortex is one of the most significant parts of the human brain and inextricably links to many advanced capacities such as complex language, social interaction, and emotional processing^[5]^. The cerebral cortex is a multilayered neuronal tissue with an interlayer distance of around 400 μm^[6]^. In addition, the cortical column is a vertical cytoarchitecture that extends through the entire cerebral cortex and has a diameter of 500 μm^[7]^. Therefore, constructing biological products with multiscale cytoarchitectures is critical for recapitulating their specific functions.

Bioprinting is one of the major biofabrication technical routes that has the ability to deposit cell-containing bioinks in a spatially-controlled manner, thus fabricating biological products with biomimetic and multiscale cellular structures^[8–11]^. By integrating customized coaxial nozzles into an extrusion bioprinting system, various multiscale heterocellular structures such as the spinal cord, hepatic lobule, and capillary were fabricated^[12, 13]^. As another bioprinting technique, digital light processing (DLP) bioprinting offers the flexibility to adjust the geometry and size of the aim tissue constructs by dynamically changing the digital masks^[14]^. In addition to bioprinting, bioassembly is recently regarded as a critical alternative by the International Society of Biofabrication^[15]^. Since the bioassembly technique can directly manipulate millions of live cells to form multicellular structures with close intercellular proximity, it improves contact-dependent cell communication and promotes the emergence of tissue-specific functions^[16–18]^. Currently, a variety of bioassembly techniques have been demonstrated in tissue construction by exploring the interactions between force fields and live cells such as electric field^[19]^, magnetic field^[20, 21]^, and acoustic field^[22–24]^. Particularly, several acoustic bioassembly techniques based on Faraday wave^[25, 26]^, bulk acoustic wave^[27–29]^, and surface acoustic wave^[30, 31]^ have been increasingly reported in biofabrication due to their advantages of high tunability, biocompatibility, and efficiency^[24]^. By using these emerging acoustic bioassembly techniques, especially the acoustic holography technique, tissues that have complex and even arbitrary cellular structures are expected to be fabricated^[32–35]^. Furthermore, the differential acoustic bioassembly technique has been demonstrated to simultaneously assemble heterogeneous cells to construct heterocellular structures based on the differences in the inherent physical properties (size and buoyant density) of the cell-containing building blocks^[36]^. While acoustic bioassembly has been rapidly advanced, the current techniques are limited to generating cytoarchitecture with a single characteristic scale that is determined by half acoustic wavelength. Therefore, it remains a great challenge for acoustic bioassembly to reconstitute multiscale cellular structures in native tissues.

To address this challenge, herein we report an emerging acoustic bioassembly technique that employs multifrequency control of Faraday wave to form multiscale cellular structures. A variety of novel assembly patterns with multiscale characteristics were obtained by using the multifrequency-driven Faraday wave bioassembly technique. Meanwhile, we conduct finite element numerical simulations to predict the spatial distributions of cell-containing building blocks in the acoustic field. We utilize two-frequency-driven Faraday wave bioassembly (f_1_=65 Hz, f_2_=148 Hz) to construct multiscale neuronal networks. hPSC-derived neural stem cells (NSCs) were assembled in Matrigel, followed by 14-day and 21-day neuronal differentiation and development. The results of immunofluorescence staining and neuroelectrophysiological analysis showed that the models have brain-specific neural types and neuroelectrophysiological functions.

## 2. Materials and methods

### 2.1. Experimental setup

The experimental setup for multifrequency-driven acoustic bioassembly was illustrated in Figure S1. The vertical vibration exciter (SA-JZ005T, Shiao, Wuxi, China) was electrically driven by an arbitrary function generator (AFG3052C, Tektronix, OR, USA) and a power amplifier (DTA-120, Dayton Audio, OH, USA). Homemade circular chambers for multifrequency-driven acoustic bioassembly were fabricated from polymethylmethacrylate (PMMA) by using a water-cooling CO_2_ laser cutting system (350-50 W, Longtai laser, Liaocheng, China). The chamber was connected with the vertical vibration exciter via a screw. The assembly chamber was 20 mm in diameter and 1.5 mm in height. In addition, confocal dishes (D29-14-1.5P, Kawei, Shanghai, China) were used for neuronal network construction. The circular chamber at the bottom of the confocal dish was 14 mm in diameter and 1 mm in height. The confocal dish was fixed on the vertical vibration exciter by using a double-sided adhesive. The level of the assembly chamber and the confocal dish were adjusted by using a bubble level.

### 2.2. PS microsphere assembly

PS microspheres (Cospheric, CA, USA) with a diameter of 100 μm were used as alternatives to living cell spheroids in the preliminary experiments to determine the multiscale effect in multifrequency-driven Faraday wave bioassembly. Specifically, microsphere suspensions with a concentration of 30 mg/mL were prepared in 1× PBS (Thermo Fisher Scientific, MA, USA) with 0.05% Tween 20 (Sigma, MO, USA) and added to the assembly chamber. Single-frequency-driven or multifrequency-driven Faraday wave was applied for microsphere assembly. A single-lens reflex camera (D3400, Nikon, Japan) was used for image acquisition.

### 2.3. Finite element analysis of Faraday wave

Finite element analysis was conducted to characterize the assembly of NSC spheroids under Faraday wave. The Faraday wave was caused by body force *F* and generated at the gas-liquid interface with a fixed boundary in a circular assembly chamber. The body force *F* was generated by a sinusoidal acceleration signal applied to the vertical vibration exciter, which can be written as:

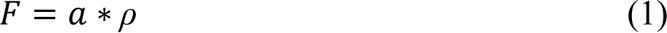

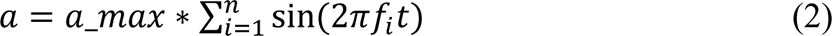

In which, *ρ* is the density of the fluid, *a_max* is the maximum acceleration applied to the vertical vibration exciter, and *f_i_*is the predefined frequency to the exciter. The Faraday wave can be simulated by applying this external force when solving the governing equation of the dynamic flow motion. These governing equations refer to the Navier-Stokes equations, which can be written as:

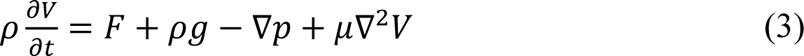

In which, *V* is the velocity vector of the flow field, *F* is the body force in Equation (1), *μ* is the dynamic viscosity of fluid, and *g* is the acceleration of gravity, respectively. For a cell spheroid at the bottom of the assembly chamber under the Faraday wave, three component forces are acting on the cell spheroid, including gravity, buoyancy and Stokes drag force. However, since gravity and buoyancy are constant values, which are only determined by the physical properties of microsphere and liquid, the position of cell spheroid is manipulated by Stokes drag force *F_drag_*. The time-average *F_drag_* as a gradient of a potential function *U* can be written as:

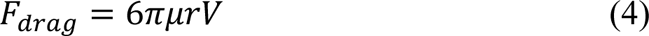

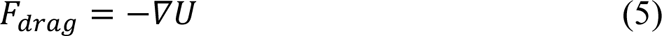

In which, *r* is the radius of the cell spheroid.

All of the simulations were performed in COMSOL Multiphysics. To reduce the computational cost, we built a simplified mathematical physics model. “Laminar Flow” physics was used to solve the governing equations of dynamic fluid motion, and the deformation of fluid motion was handled by adding a “dynamic mesh” condition. The “volume force” condition was applied to the 3D calculation domain of “Laminar Flow” physics. The boundary conditions on the gas-liquid interface and solid-liquid interfaces were set to “free surface” (surface tension of gas-liquid interface *σ*) and “no slip”, respectively. The problem was solved via a “Time-Dependent” solver. The parameters of materials used in the calculations are given in Table 1.

**Table 1.** The parameters of materials.

|  |  |  |
| --- | --- | --- |
| Dynamic viscosity of fluid | $\mu$ | $1.01 \times 10^{-3} \text{ Pa} \cdot \text{s}$ |
| Surface tension of gas-liquid interface | $\sigma$ | 45 mN/m |
| Maximum acceleration applied to the exciter | $a_{max}$ | $2 \cdot g$ |
| Acceleration of gravity | $g$ | $9.80 \text{ m/s}^2$ |
| Radius of cell spheroid | $r$ | 125 $\mu\text{m}$ |

### 2.4. hNSC differentiation

The hPSCs were cultured in NuwacellTM ncTarget hPSC Medium (Nuwacell Biotechnologies, RP01020, Hefei, China) with 10 μM Y-27632 (STEMCELL Technologies, 72304, BC, USA). The hNSCs were derived from hiPSCs which followed standard neurobiology protocols (Thermo Fisher Scientific). Specifically, the hiPSCs were cultured in NuwacellTM ncTarget hPSC Medium on day 1 of splitting with 25% confluency. Then the cells were subsequently cultured in neural induction medium containing neurobasal medium and neural induction supplement. On day 7 of neural induction, hNSCs derived from hPSCs were ready to be harvested and expanded.

### 2.5. hNSC spheroids fabrication

hNSCs were seeded into a 6-well ultralow adhesion microtiter plate (Corning, 3471, NY, USA) at 4 × 10^5^ cells/well. The plate was placed on an orbital shaker (GS-10, MiuLab, Hangzhou, China) at the speed of 75 rpm for 1 day for the formation of hNSC spheroids. hNSC spheroid formation and culture were both maintained at 37℃ with 5% CO_2_. The formed hNSC spheroids were collected and utilized in Faraday wave bioassembly.

### 2.6. Neuronal network construction

hNSC spheroids were uniformly resuspended in 4∼6 mg/mL Matrigel (Corning, 356255, AZ, USA) and loaded into the chamber of a confocal dish. After the sedimentation of hNSC spheroids, a two-frequency-driven Faraday wave was applied for the generation of the predefined cellular structure (f_1_=65 Hz, f_2_=148 Hz). After that, the tissue constructs were transferred to an incubator for 30 minutes for gelation and immobilization. The tissue constructs were firstly cultured in Neural Stem Cell Serum Free Medium (Thermo Fisher Scientific, A1050901, MA, USA) for 1 day and then differentiated in neural differentiation medium for 13 days to obtain neuronal network models.

### 2.7. Cell viability assay

The cell viability was examined by using Calcein-AM/PI staining. Tissue constructs were washed with 1× PBS 3 times and incubated for 30 min at 37 °C in 1 mL staining solution containing 0.1% (v/v) Calcein-AM solution and 0.1% (v/v) PI solution (Beyotime Biotechnology, Shanghai, China). After that, tissue constructs were imaged under an inverted fluorescence microscope (IX83, Olympus, Japan). ImageJ software (NIH, Bethesda, MD, USA) was used to count the live and dead cells for each image. The cell viability was calculated as below,

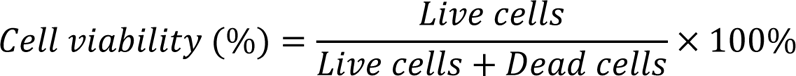

### 2.8. Immunostaining

Neural progenitors and neurons in the formed neuronal networks were analyzed by Immunostaining. Specifically, neuronal networks were fixed in 4% paraformaldehyde (Sigma, MO, USA) for 30 minutes at room temperature. The samples were washed with 1× PBS supplemented with 0.5% Triton X-100 (Sinopharm, Beijing, China) 3 times. And then the samples were blocked with 1% bovine serum albumin (BSA) (Sigma, MO, USA) supplemented with 0.1% saponin (Sigma, MO, USA) overnight at 4℃. After that, the samples were incubated with primary antibodies in the diluent solution at 4℃ for 48 h and then washed with 1× PBS supplemented with 0.5% Triton X-100 3 times on a rocker. Then, the samples were incubated with secondary antibody in the diluent solution at 4 °C for 24 h in the dark, and washed with 1× PBS supplemented with 0.5% Triton X-100 3 times on a rocker. Finally, nuclei were counterstained with DAPI (Thermo Fisher Scientific, MA, USA) for 30 minutes. The images were acquired using laser scanning confocal microscopy (SP8 STED, Leica, Germany).

### 2.9. Neuroelectrophysiological Measurement

The neuroelectrophysiological function of the formed neuronal networks was assessed by sampling spontaneous electrical activity 14 days after hNSC assembly. Briefly, the formed neuronal networks in 6-well plates integrated with MEA (Axion BioSystems, Maestro Pro) were placed in a carbon dioxide-flooded measurement device. Data were collected using an AxIS Navigator (Axion BioSystems) and exported to NeuralMetric Tool, and the traces were processed to extract spontaneous spikes.

### 2.10. Data analysis

The data were plotted by using GraphPad Prism 8 (GraphPad Software, San Diego, CA, USA) software. All data are presented as mean ± SD values and were analyzed by SPSS Statistics 21 software. Differences were considered significant at *P < 0.05, **P < 0.01, and ***P < 0.001.

## 3. Results and discussion

In nature, any periodic wave can be decomposed into a series of superpositions of sine waves and cosine waves through the Fourier transform (Figure 1A). Typically, a single-frequency-driven Faraday wave has a unique waveform pattern which is determined by the mode number pair (m, n). Specifically, the m and n present the number of nodal diameters and nodal circles, respectively^[37, 38]^. During the bioassembly process, the waveform pattern can be transferred from the air-liquid interface to building blocks in the acoustic field. In this study, we superimposed multiple sine wave signals to induce Faraday waves with different modes concurrently existed and consequently obtained the assembled structures that have much richer variations (Figure 1B)^[39]^.

**Figure 1.**
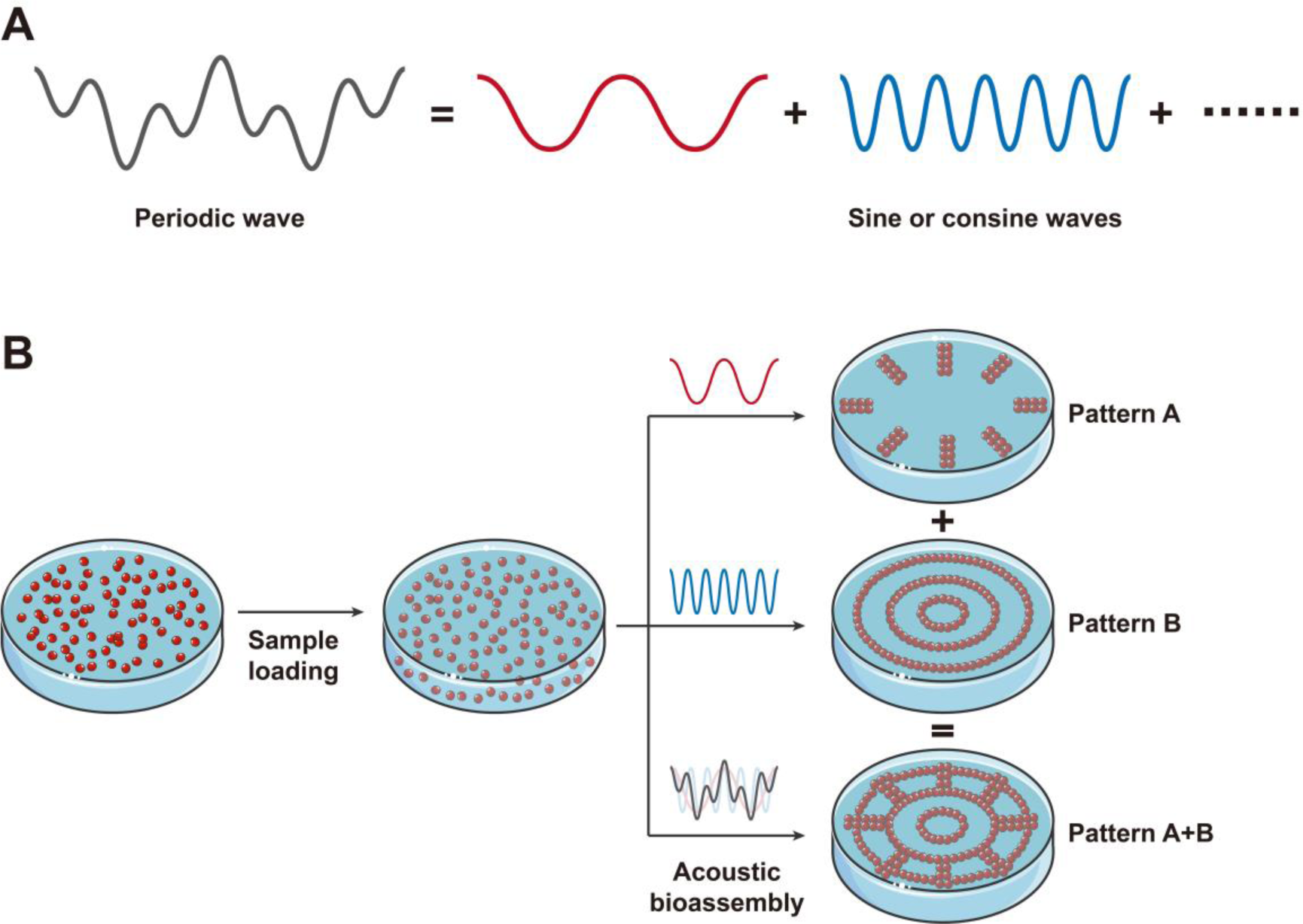
Scheme of the multifrequency-driven Faraday wave bioassembly. (A) A periodic wave can be decomposed into a sum of sine or cosine waves through Fourier series. (B) The procedure of the multifrequency-driven Faraday wave bioassembly.

To investigate the law of the multifrequency-driven Faraday wave bioassembly, we first employed a series of single-frequency signals to induce the Faraday wave. We used hNSC spheroids as building blocks and obtained 15 nodal patterns by increasing the driving frequency from 15 Hz to 100 Hz (Figure 2A). We observed that the number of nodal diameters and nodal circles of the assembly patterns showed an increasing tendency as the driving frequency increased (Figure 2B-C). The patterns in the same line had the same number of nodal circles. Similarly, the number of nodal diameters for the assembly patterns in each column was equal (Figure 2A). Next, we superimposed two sine wave signals to induce the Faraday waves, where the signal function can be written as *y* = *A*_1_sin(35 × 2*πt*) + *A*_2_sin (74 × 2*πt*) ^[40]^. In the classic single-frequency-driven Faraday wave bioassembly, the Faraday wave was generated as soon as the vertical mechanical vibration of the liquid layer in the assembly chamber exceeded an acceleration threshold that induced hydrodynamic instability of the gas-liquid interface^[41]^. When the amplitude ratio (*A*_1_/*A*_2_) between two sine wave signals was equal to 0.8, we obtained a concentric circle pattern due to the Faraday wave with 74 Hz driving frequency first reached and exceeded the acceleration threshold. Based on the same principle, a radial pattern was generated when the amplitude ratio was equal to 1.4. Interestingly, Faraday waves with 35 Hz and 74 Hz driving frequencies almost exceeded the acceleration threshold simultaneously and kept in balance when the amplitude ratio was set at 1.0 or 1.2. As a result, a new assembly pattern was generated that totally contained concentric circles and radial features (Figure 2D). Further, the sine wave signal in 74 Hz was selected to separately superimpose with other 14 signals to assemble hNSC spheroids under two-frequency-driven Faraday wave bioassembly. As shown in Figure 2E, the generation of new assembly patterns was strictly dependent on the number of nodal diameters and nodal circles of the original single-frequency-driven patterns. In the first column, the three-concentric-circle pattern contained all the features of the other two patterns because its number of nodal circles was larger. Similarly, the other four patterns in the last line had a larger number of diameters, thus they had all the features of the three-concentric-circle pattern. Overall, two-frequency-driven assembly patterns with more complex features could be obtained just when the number of nodal diameters and nodal circles were both different in the single-frequency-driven patterns.

**Figure 2.**
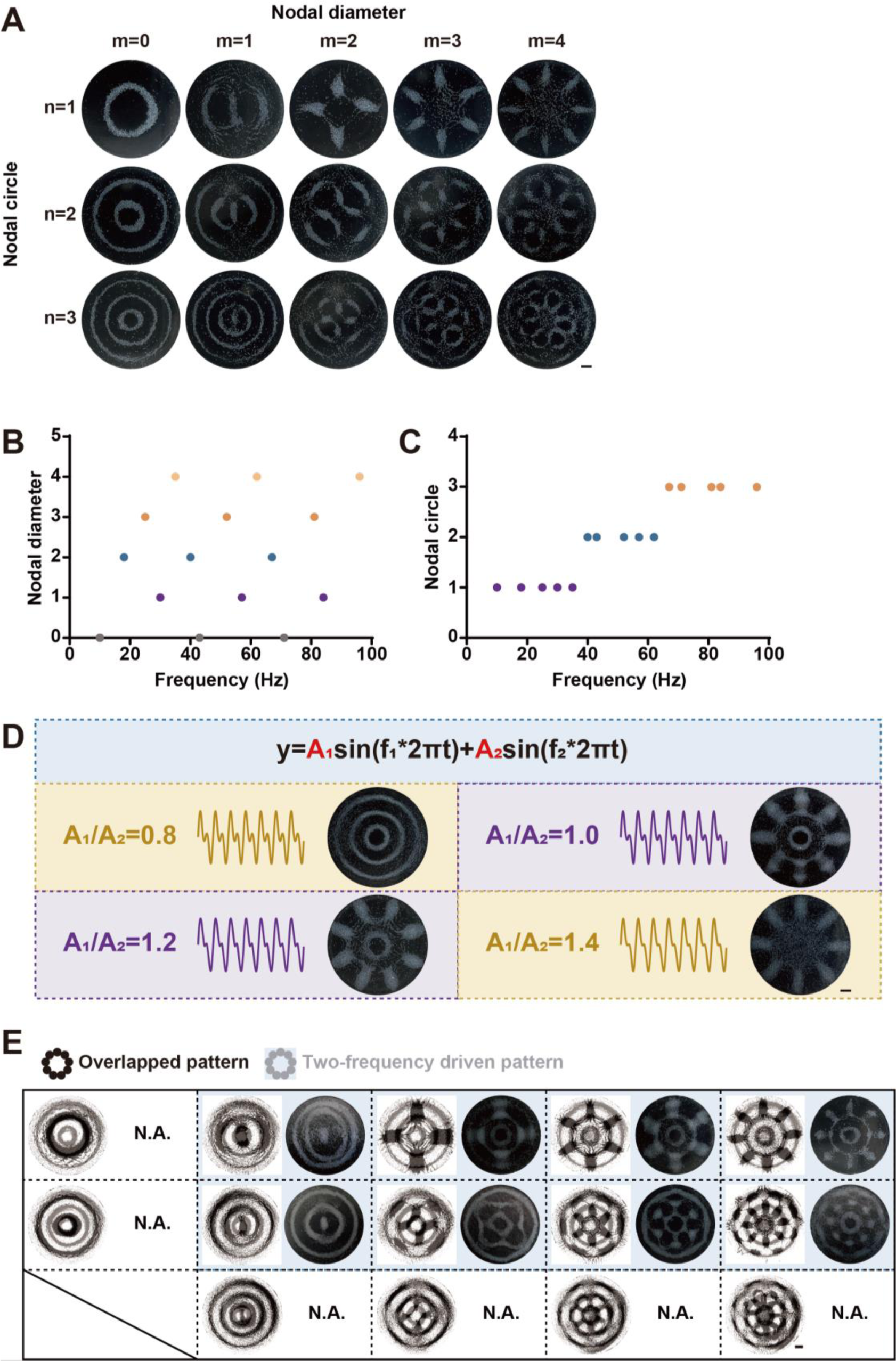
The law of the multifrequency-driven Faraday wave bioassembly. (A) Nodal assembly patterns formed by hNSC spheroids. (B) The relationship between the number of nodal diameter and the driving frequency of the Faraday wave. (C) The relationship between the number of nodal circle and the driving frequency of the Faraday wave. (D) Two-frequency-driven Faraday wave bioassembly by tuning the amplitude ratio between two sine wave signals. (E) The influence of the number of nodal diameter and nodal circle on multifrequency-driven Faraday wave bioassembly. All scale bars indicate 2 mm.

Using the multifrequency-driven Faraday wave bioassembly technique, we obtained several two or three-frequency-driven patterns. Based on the established mathematical physical model, we numerically simulated the force potential distribution as well as the streamlines of the flow field, which can be used to accurately predict the aggregation regions of hNSC spheroids (Figure 3A and S2). The regions with the lowest force potential marked in dark blue were consistent with the actual aggregation area of hNSC spheroids (Figure 3A). To further investigate the structural characteristics of multifrequency-driven patterns, we quantitatively analyzed the changes in their number of nodal circles and nodal diameters as well as the distributions of the hNSC spheroids. We observed that the number of nodal circles and nodal diameters in multifrequency-driven patterns equaled the maximum value in the corresponding single-frequency-driven patterns (Figure 3B). Meanwhile, the multifrequency-driven patterns could obtain all the features of the corresponding single-frequency-driven patterns (Figure 3C). Together, more complex patterns that inherited all the original structural features could be generated by using the multifrequency-driven Faraday wave bioassembly.

**Figure 3.**
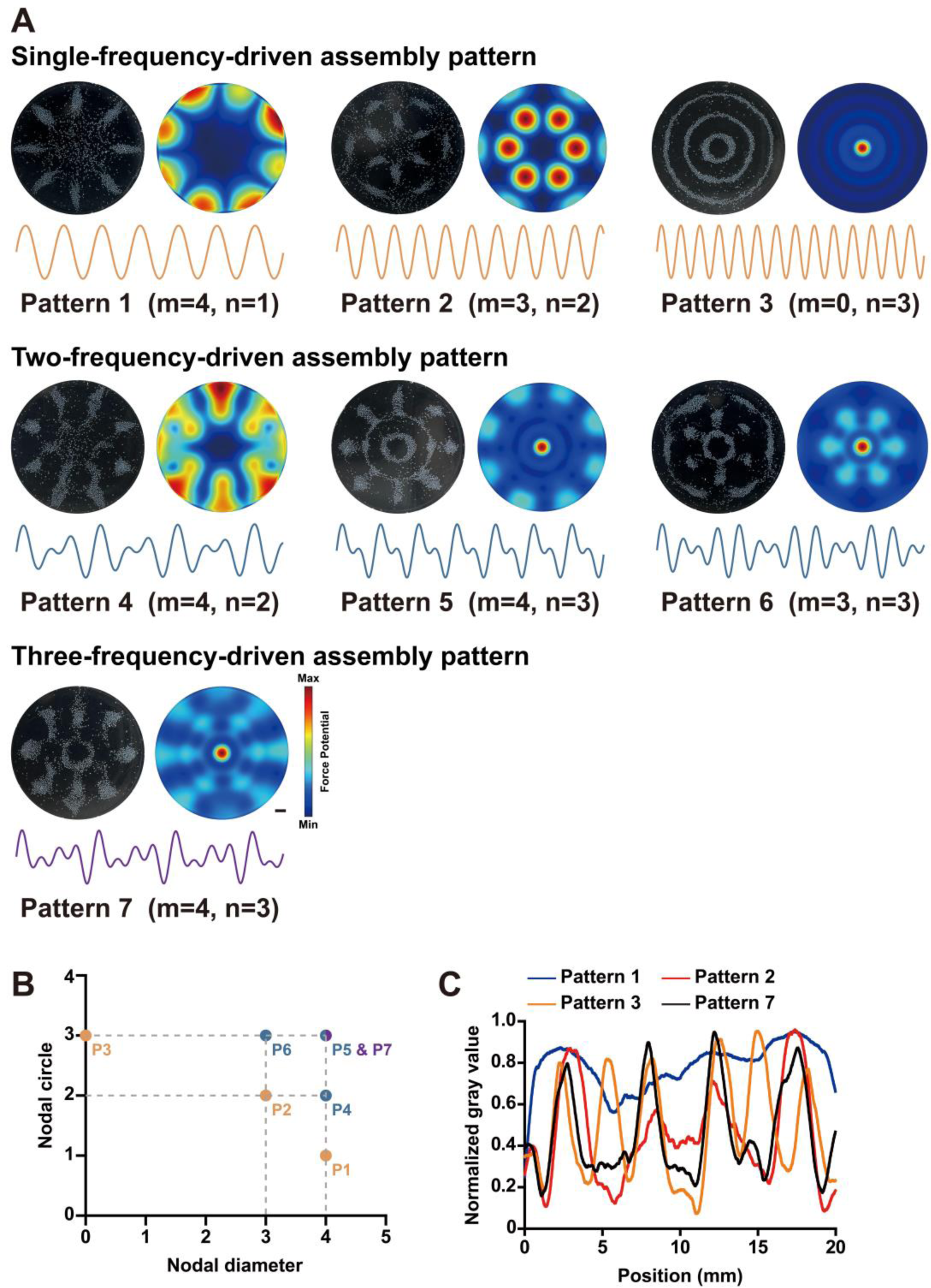
Multifrequency-driven Faraday wave bioassembly. (A) Assembly patterns fabricated by single, two or three-frequency-driven Faraday wave bioassembly. (B) The change rule of the nodal diameters and nodal circles in multifrequency-driven patterns. (C) Quantitative analysis the distributions of the single-frequency-driven patterns and three-frequency-driven pattern. Scale bar: 2 mm.

In acoustic bioassembly, the building blocks will aggregate at the nodes or antinodes of the acoustic waves. Thus, the characteristic scale of the assembly patterns was determined by the half-wavelength of the acoustic waves^[42]^. The working frequency of the Faraday wave, bulk acoustic wave, and surface acoustic wave increases in turn, from hectohertz to gigahertz^[43–46]^. Based on the higher working frequency compared with the Faraday wave, the bulk acoustic wave and surface acoustic wave have a strong ability to assemble single cells into specific cellular structures with smaller characteristic scales. However, the working frequency of a piezoelectric transducer used for bulk acoustic wave generation covers only a bandwidth of a few hectohertz^[47, 48]^. Moreover, a piezoelectric transducer can just work with a single frequency at a time. And for surface acoustic wave, the distance between two adjacent interdigital electrodes determines the frequency of the wave^[49]^. The working frequency of a surface acoustic wave cannot be changed as soon as the interdigital transducer is fabricated. Based on the reasons aforementioned, it is difficult to induce acoustic waves with multiple work frequencies in a single bioassembly process by employing piezoelectric transducers or interdigital transducers. In multifrequency-driven Faraday wave bioassembly, multiple Faraday waves can be induced simultaneously with distinct half-wavelength, and thus meet the requirement for constructing multiscale tissue mimics. The half wavelength of the Faraday wave can be expressed as^[50]^:

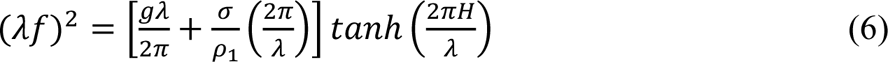

Where *f* is the working frequency of the Faraday wave, *λ* is the wavelength of Faraday waves, *g* is the gravitational acceleration, *σ* is the surface tension of the liquid, *ρ*_1_ is the liquid density, and *H* is the height of the assembly chamber.

In a circular chamber with 6 cm in diameter and 1.5 mm in height, we assembled single-frequency-driven patterns by tuning the driving frequency from 20 Hz to 168 Hz (Figure 4A). We compared the characteristic scale of these assembly patterns with the theoretical value (Figure 4B-C). As the driving frequency of the Faraday wave increased, the characteristic scale of assembly patterns gradually decreased and they well matched the theoretical value of the Faraday wave half-wavelength. Further superimposed two frequency signals, assembly patterns with multiscale features were generated (Figure 4D). Taken together, the multifrequency-driven acoustic bioassembly technique can meet the requirement for the construction of multiscale cellular structures. By using equation (6), we can also rationally design tissue mimics with predefined scales. However, in this study, the minimum features of assembly patterns were on a millimeter scale which could not reconstitute micron-scale cellular structures in native tissues. We expect to reduce the minimum feature size by employing a high-frequency vibrational exciter.

**Figure 4.**
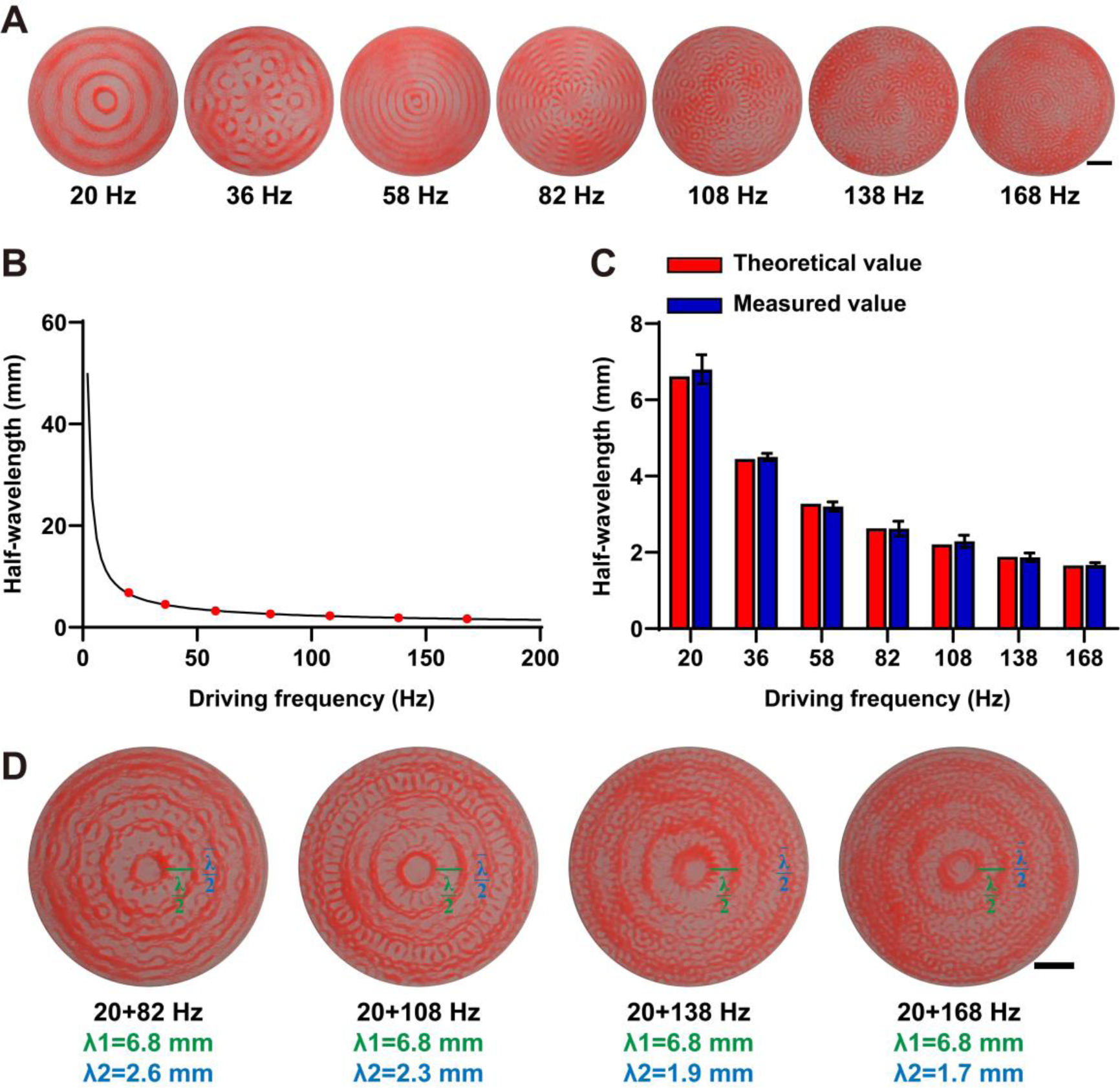
Assembly of patterns with multiscale structures. (A) Single-frequency-driven assembly patterns. (B) The relationship between the half-wavelength and the driving frequency of the Faraday wave. (C) The comparison between the theoretical half-wave length of the Faraday wave and the characteristic scale of the single-frequency-driven assembly patterns. (D) Two-frequency-driven assembly patterns with two characteristic scales. All scale bars indicate 10 mm.

As a proof-of-concept work, we constructed neuronal network models using a multifrequency-driven acoustic bioassembly technique to validate its potential in fabricating tissue mimics with multiscale cellular structures. hNSC spheroids with 125.7 µm were used as building blocks for neuronal network model construction (Figure S3A-C). We investigated the biocompatibility of the multifrequency-driven acoustic bioassembly technique. Specifically, the live cells were stained with Calcein-AM in green fluorescence and dead cells were stained with PI in red fluorescence (Figure S3D). The cell viability of hNSCs before and after bioassembly was 88.4% and 90.4%, respectively. The result revealed that the cell viability was not affected by the bioassembly process (Figure S3E). During the culture process, hNSC spheroids fused over time and the assembled cellular structure became an integrated construct without losing the original patterns (Figure 5A). Furthermore, we identified the cell populations and neuroelectrophysiological functions of neuronal networks. Immunofluorescence staining demonstrated that neurons in the assembled constructs formed multiscale synaptic connectivity and expressed TuJ1 (early-stage neuronal biomarker) and MAP2 (post-mitotic mature neuron-specific biomarker) (Figure 5B and S4). Neuroelectrophysiological analysis revealed that the neuronal network models could spontaneously generate neuroelectrophysiological activities (Figure 5C). Specifically, the mean firing rate was 0.16 Hz, the number of spikes in a minute was 580, and the number of bursts in a minute was 16 (Figure 5D-F).

**Figure 5.**
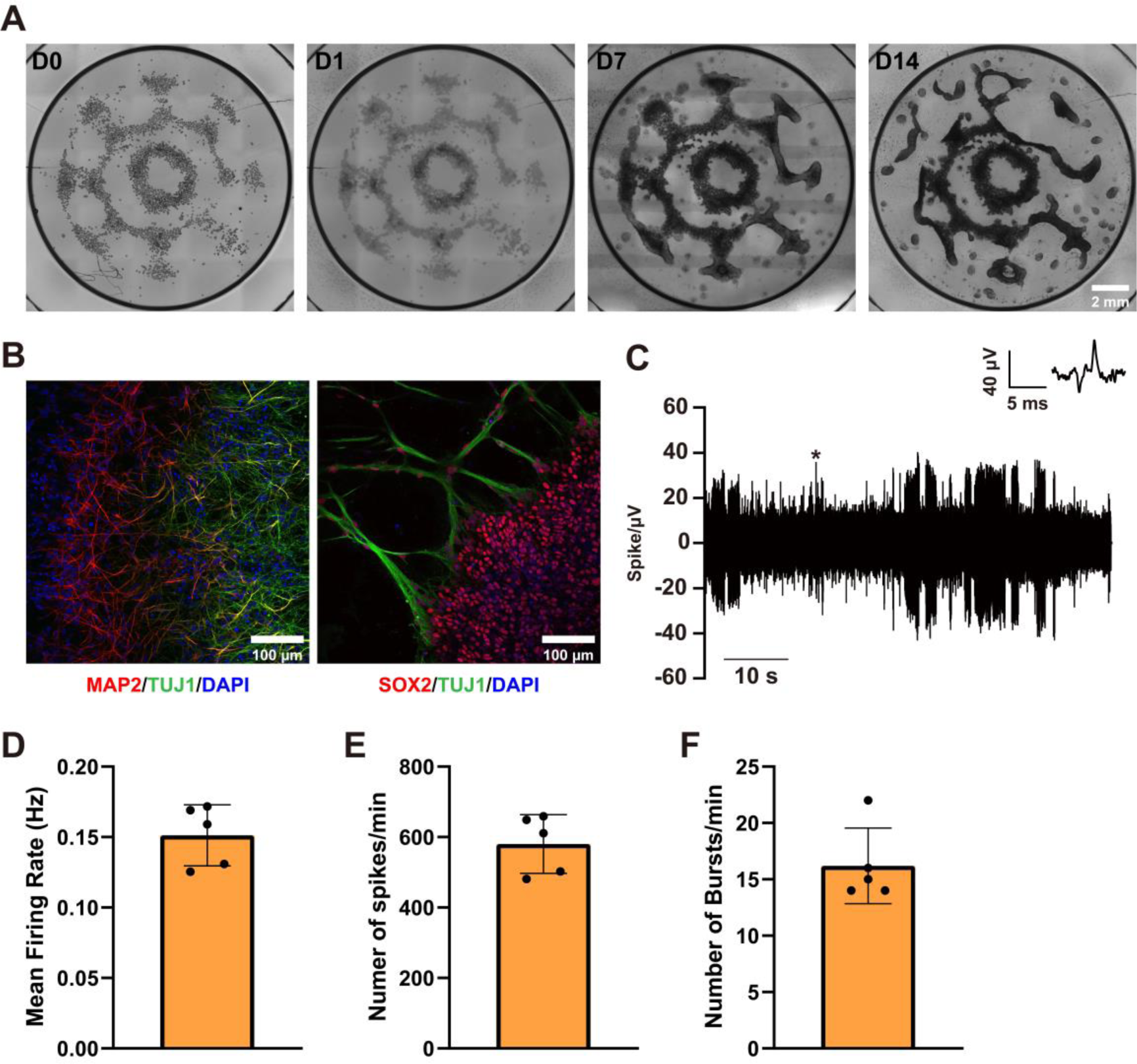
Construction and function evaluation of neuronal network models. (A) In vitro culture of neuronal network models. (B) Immunofluorescence staining of neuronal network models with TUJ1, MAP2 and SOX2. (C) Relative voltage changes in neuronal network models. (D) Mean firing rate in neuronal network models. € Number of spikes in neuronal network models. (F) Number of bursts in neuronal network models.

## 4. Conclusion

In this work, we reported a novel acoustic bioassembly technique that employed multifrequency control of Faraday waves to form multiscale cellular structures. This technique overcomes the structural limit of single-frequency driven acoustic bioassembly and potentially emulates complex cytoarchitecture in natural tissues. Additionally, this technique preserves the traditional advantages of bioassembly and enables to generate multiscale structures in a few seconds. Using this technique, we constructed functional neuronal networks with multiscale connectivity that displayed electrical activities. We expect this technique will find wide applications in tissue engineering and regenerative medicine.

## Acknowledgments

The authors gratefully acknowledge the financial support from the National Natural Science Foundation of China (Grant No: 82272173).

## Declaration of competing interest

P.C. is a cofounder of, and has an equity interest in: (i) Shenzhen Convergence Bio-Manufacturing Co., Ltd., a company that is developing convergence bio-manufacturing technologies to enable artificial meat and regenerative medicine. P.C.’s interests were viewed and managed in accordance with the conflict of interest policies.

## CRediT authorship contribution statement

**Longjun Gu:** Data curation, Formal analysis, Investigation, Methodology, Software, Validation, Visualization, Writing - original draft, Writing - review & editing. **Zhao Wen:** Formal analysis, Investigation, Methodology. **Yuhang Fan:** Data curation, Formal analysis, Investigation, Validation, Visualization. **Jia Shang:** Investigation, Methodology. **Yang Zhao:** Investigation, Validation. **Jibo Wang:** Software. **Tao Chen:** Investigation. **Peidi Liu:** Visualization. **Pu Chen:** Conceptualization, Data curation, Funding acquisition, Investigation, Methodology, Project administration, Resources, Software, Supervision, Validation, Writing - original draft, Writing - review & editing.

