## Supplementary Information for "Multifrequency control of Faraday wave bioassembly for constructing multiscale hPSC-derived neuronal networks"

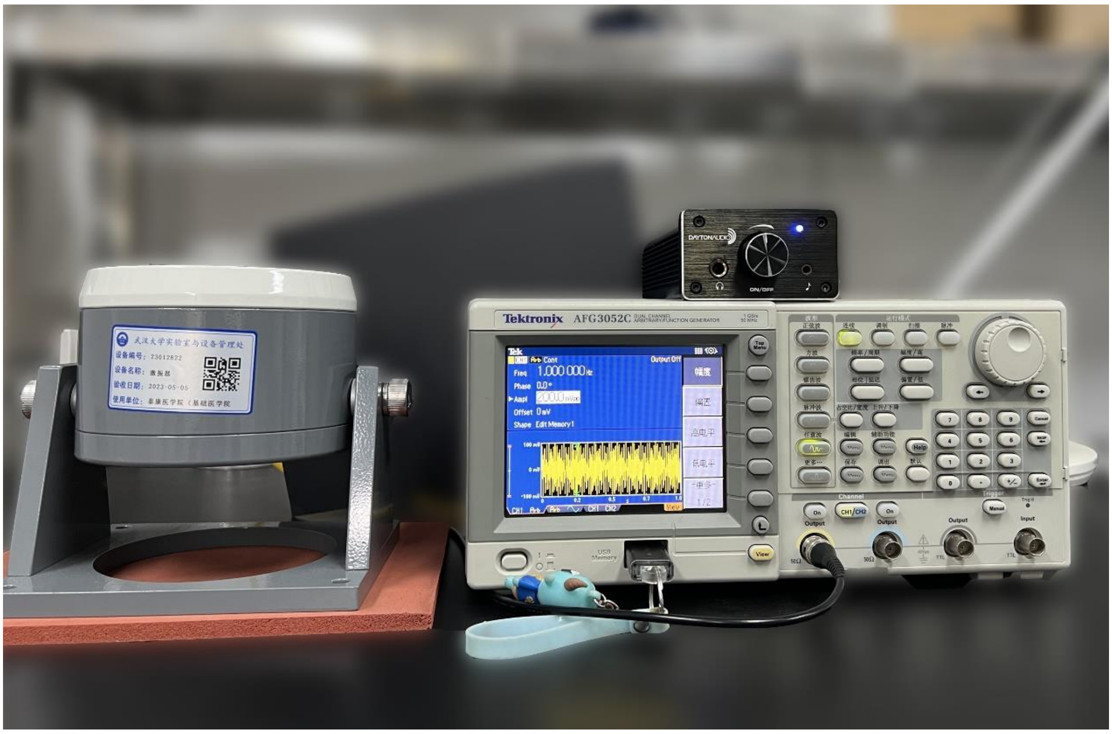


Figure S1. Experimental apparatus for multifrequency-driven Faraday wave bioassembly technique.

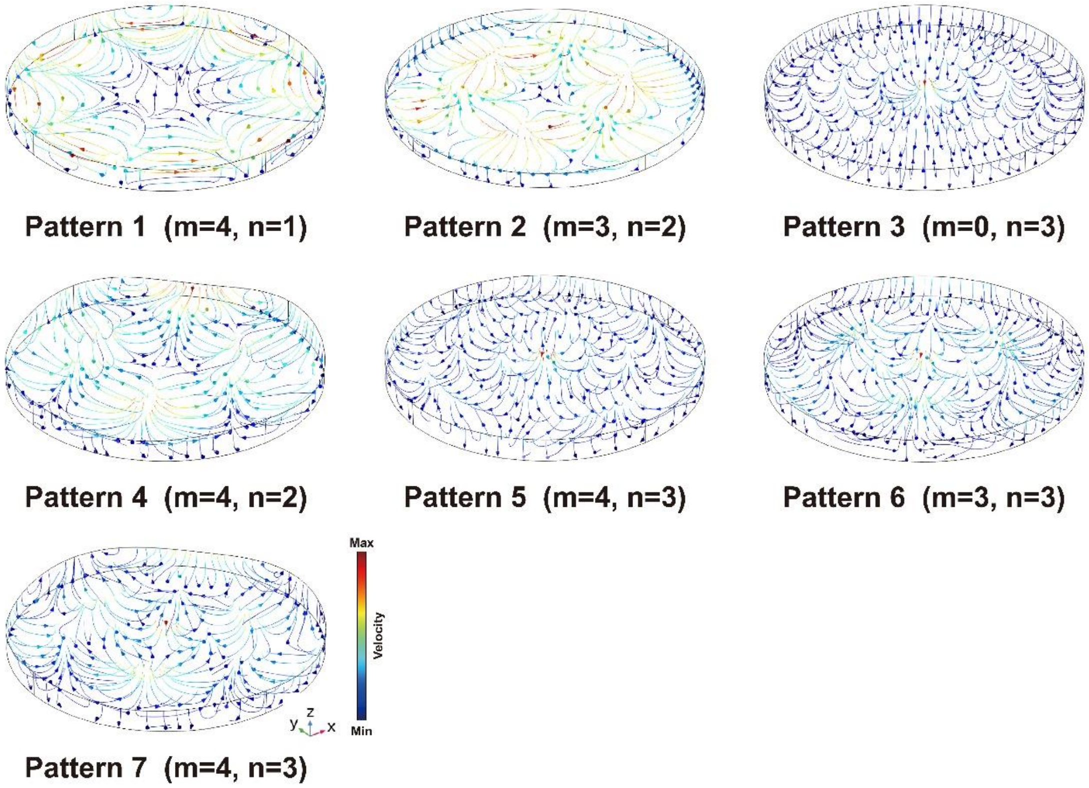


Figure S2. The finite element numerical simulation of the streamlines in the flow field. Pattern 1 to pattern 3 are generated by single-frequency-driven Faraday wave bioassembly. Pattern 4 to pattern 6 are generated by two-frequency-driven Faraday wave bioassembly. Pattern 7 is generated by three-frequency-driven Faraday wave bioassembly.

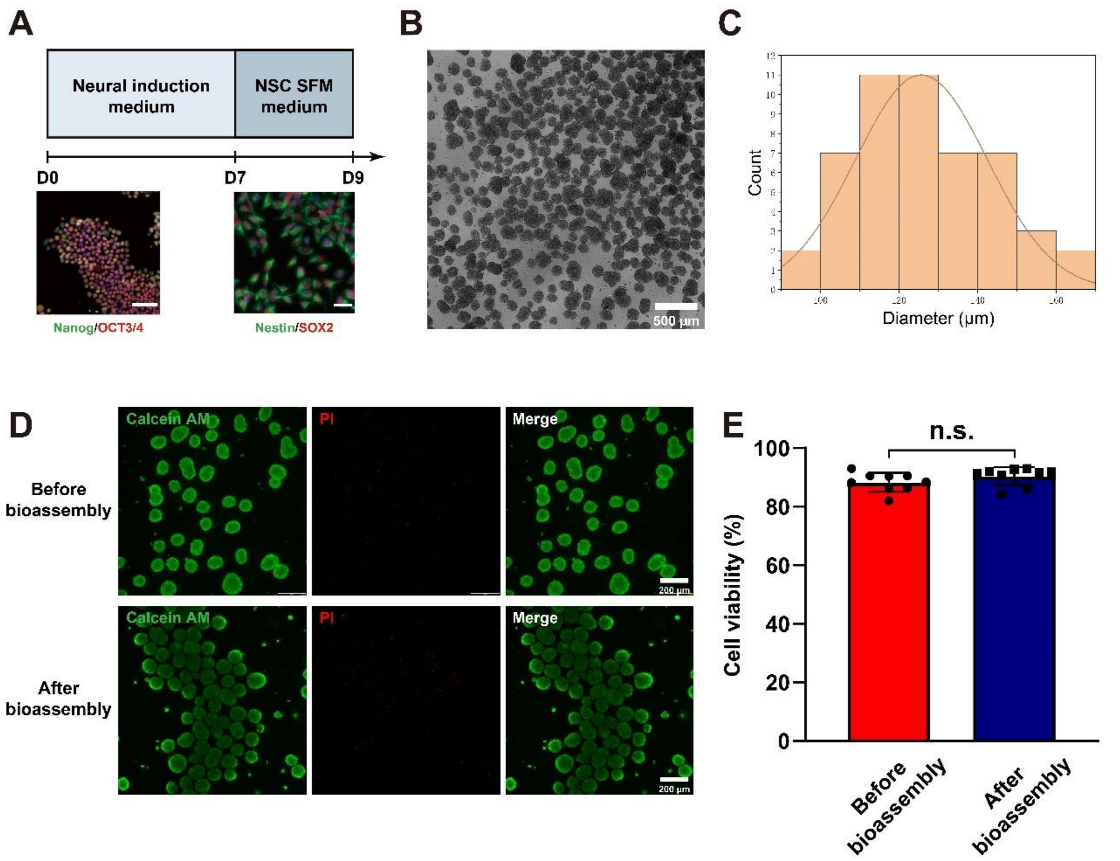


Figure S3. Fabrication and evaluation the cell viability of hNSC spheroids. (A) Differentiation process of hNSCs. hiPSCs were positively marked with Nanog and OCT3/4. hNSCs were positively marked with Nestin and SOX2. Scale bar: 100μm. (B) The bright field image of the formed hNSC spheroids. (C) Size distribution of the hNSC spheroids. (D) Evaluation of cell viability by Calcein AM/PI assay before and after acoustic bioassembly. (E) Quantitative statistics of the cell viability.

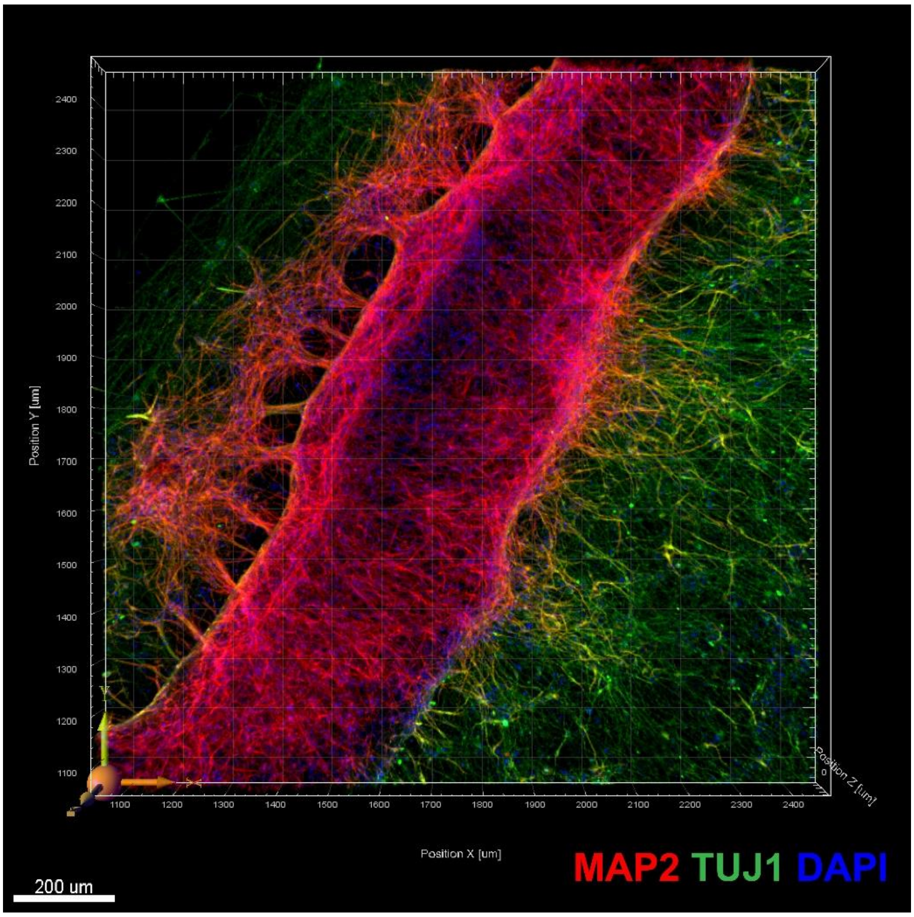


Figure S4. Fluorescent images of a part of the neuronal network models.
